# Posterior insula oscillations are not modulated by changes in perceived intensity: Evidence from human intracerebral EEG

**DOI:** 10.64898/2026.07.17.739090

**Authors:** Chiara Leu, Aïcha Boutachkourt, Susana Ferrao Santos, Alexane Fierain, Vincent Joris, Patrice Finet, Giulia Liberati

## Abstract

The human insular cortex is involved in a multitude of processes essential for survival, but its precise role in pain perception remains debated. While it has been shown that ongoing oscillatory activity in the posterior insula is preferentially modulated by thermonociceptive input, it is unclear whether these modulations are functionally related to changes in pain perception. To assess this link, neural responses to sustained periodic thermonociceptive and non-nociceptive vibrotactile stimulation delivered at a frequency of 0.2 Hz were measured using intracerebral electrode contacts located in the anterior (n=63) and posterior (n=37) insular cortices of 10 patients undergoing presurgical evaluation for focal epilepsy. An arithmetic task was employed concomitant to the stimulation, aiming to reduce participant’s perceived intensity while leaving the physical stimulation parameters unchanged. A frequency-tagging analysis approach was used to assess the modulation of the ongoing neural oscillations. As expected, the perception of the stimulation was significantly reduced during the arithmetic task to a similar extent in both modalities. Yet, no congruent reduction in the frequency-tagged oscillatory responses was found in the posterior insula during distraction, for either of the modalities. On the contrary, during distraction, vibrotactile stimulation elicited larger responses in the anterior insula. These findings suggest that top-down modulation via distraction differentially affects anterior insula responses to innocuous vibrotactile input, while thermonociceptive processing appears to involve additional neural systems during distraction that are not equivalently engaged by vibrotactile stimulation.

## 1. Introduction

The underlying mechanisms leading to the experience of pain caused by a nociceptive stimulus remain elusive [42]. The human insular cortex is a promising target to deepen the understanding of these mechanisms, as it has been shown to be consistently activated by nociceptive stimuli perceived as painful [2,6,21], serves as a relay and integration “hub” for sensory information, and plays an important role in interoception and emotional regulation [25,30,59,70]. Crucially, while the anterior portion of the insular cortex is mainly involved in the cognitive and emotional processing of a painful stimulus [24,26,29,61,63], some investigations argue that its activation reflects the perceived level of intensity of a nociceptive stimulus [15,17]. Conversely, activation in the posterior insula has been claimed to be predominantly related to the actual intensity of such a stimulus [23,24,57]. Overall, the distinct role of the insular cortex in the perception of pain is still disputed [16,56].

To further investigate the involvement of the insular cortex in pain perception, intracerebral electroencephalography (iEEG) is a useful tool to study the temporal aspects of pain processing with a high spatial resolution within different regions of the insula. Using a frequency-tagging approach [11,12,51], iEEG recordings from the dorsal posterior insula have shown that ongoing neural oscillations are preferentially modulated by sustained periodic thermonociceptive stimuli in the theta and alpha frequency bands at the frequency of stimulation [36]. These findings suggest a relationship between nociception and ongoing oscillations, but it remains unclear whether these modulations play a significant role in the perception of pain. If this is the case, cognitive processes known to affect pain perception should also affect these ongoing oscillations. Thus, changes in the attentional state of the subject should be reflected in changes in pain perception [32], and consequently also in the modulation of ongoing oscillations in the posterior insula. Importantly, such an attention modulation paradigm using an arithmetic distraction task has already been used in a scalp EEG investigation following the same rationale [33]. The results of this investigation were not conclusive, possibly due to the low spatial resolution of scalp EEG. It is plausible that any activity from cortical areas that are located deep in the brain would be drowned out by noise in the scalp EEG measurement [50]. Given the potential involvement of the insula in the processing of nociceptive stimuli, we expect to obtain a better understanding of the underlying mechanisms using iEEG with electrode contacts located in the anterior and posterior insula. Thus, the same ultra-slow sustained periodic stimulation paradigm at a frequency of 0.2 Hz was applied, using both thermonociceptive and vibrotactile stimulation [33].

We hypothesized that a distraction task would lead to a decrease in the perceived intensity in both modalities. A concomitant decrease in amplitude of the modulation of ongoing oscillations exerted by the stimulation would indicate a link to intensity perception. If the modulation is preferential for pain perception, the neural response following non-painful sustained periodic vibrotactile stimuli will be different compared to the thermonociceptive stimuli.

## 2. Methods

### 2.1 Participants

10 patients (age: 31.8 ± 8.7 years (mean ± standard deviation), 3 female) undergoing pre-surgical evaluation for refractory focal epilepsy were recruited at the Department of Neurology at the Saint-Luc University Hospital (Brussels, Belgium). Recruitment, electrode implantation and iEEG recordings were performed as described in our previous collaborations with the Refractory Epilepsy Centre, and our sample size matches these investigations [36–39]. The study was conducted over four years, and no further patients were enrolled beyond those available within that timeframe.

As part of their evaluation, all patients had depth electrodes implanted in different regions of the brain that were suspected to be the origin of their seizures. We focused only on electrode contacts implanted in the anterior (n=63) and posterior (n=37) insula. For 2 patients, only the thermonociceptive part of the study was conducted, due to time constraints and patient availability. Thus, for the vibrotactile modality, only 48 anterior and 24 posterior electrode contacts are included in the analysis. Insular regions were identified as either the region surrounding the short insular gyri, the pole of the insula and the transverse insular gyrus for the anterior insula or the regions encompassing the anterior and posterior long insular gyri for the posterior insula [45] (see section 2.4 for a more detailed description of contact localization). 7 participants had depth electrodes implanted in the anterior and posterior insula, 2 patients only in the anterior and 1 patient only in the posterior insula. MNI coordinates for all electrode contacts can be found in Table S.1. This data will be publicly available upon publication, containing anonymized DICOM images of each patient. For each image, it will be ensured that no facial features that could lead to the identification of the patient are recognizable.

None of the participants had a medical history (psychiatric diseases, sensory abnormalities, or cognitive deficits) that would hinder their participation in the experiment; they were therefore considered representative of a healthy population. Additionally, no ictal discharges were detected during the experiment in any of the participants.

Prior to signing an informed consent form, all participants were informed by the responsible neurologist that this experiment would not influence their treatment and that they were free to choose whether they wanted to participate. The local research ethics committee approved the investigation (Commission d’Ethique Hospitalo-Facultaire Saint-Luc UCLouvain, B403201316436).

### 2.2 Experimental setup

The same protocol as described in Leu *et al.* [33] was used for the experimental setup, carried out at the patient’s bedside. Very slow periodic sustained thermonociceptive and vibrotactile stimuli were used in separate blocks, at a frequency of 0.2 Hz over a duration of 75 s per trial, during which participants were either not requested to do anything in particular (i.e. baseline condition) or had to solve an arithmetic task (i.e. distraction task).

The temperature of the thermonociceptive stimuli varied between baseline temperature (35°C) and 50°C in a sinusoidal manner over a duration of 5 s per cycle. Stimuli were delivered using a thermal cutaneous stimulator (TCS II, QST.Lab, Strasbourg, France). The probe (model T03, set with 15 micro-Peltier elements over a 30mm diameter surface) was placed on the volar forearm of the participant, contralateral to where the highest number of insular contacts were located. The non-nociceptive vibrotactile stimuli were delivered using a 251 Hz sine wave whose amplitude was periodically modulated at a frequency of 0.2 Hz (Figure 1), using a recoil-type vibrotactile transducer (Haptuator, Tactile Labs Inc., Canada) positioned between the thumb and the index of the arm contralateral to where the highest number of insular contacts were located. This non-nociceptive stimulation, generally perceived as non-painful but intense [36], was chosen to assess the selectivity of the observed brain responses for nociceptive stimulation and/or pain perception.

**Figure 1.**
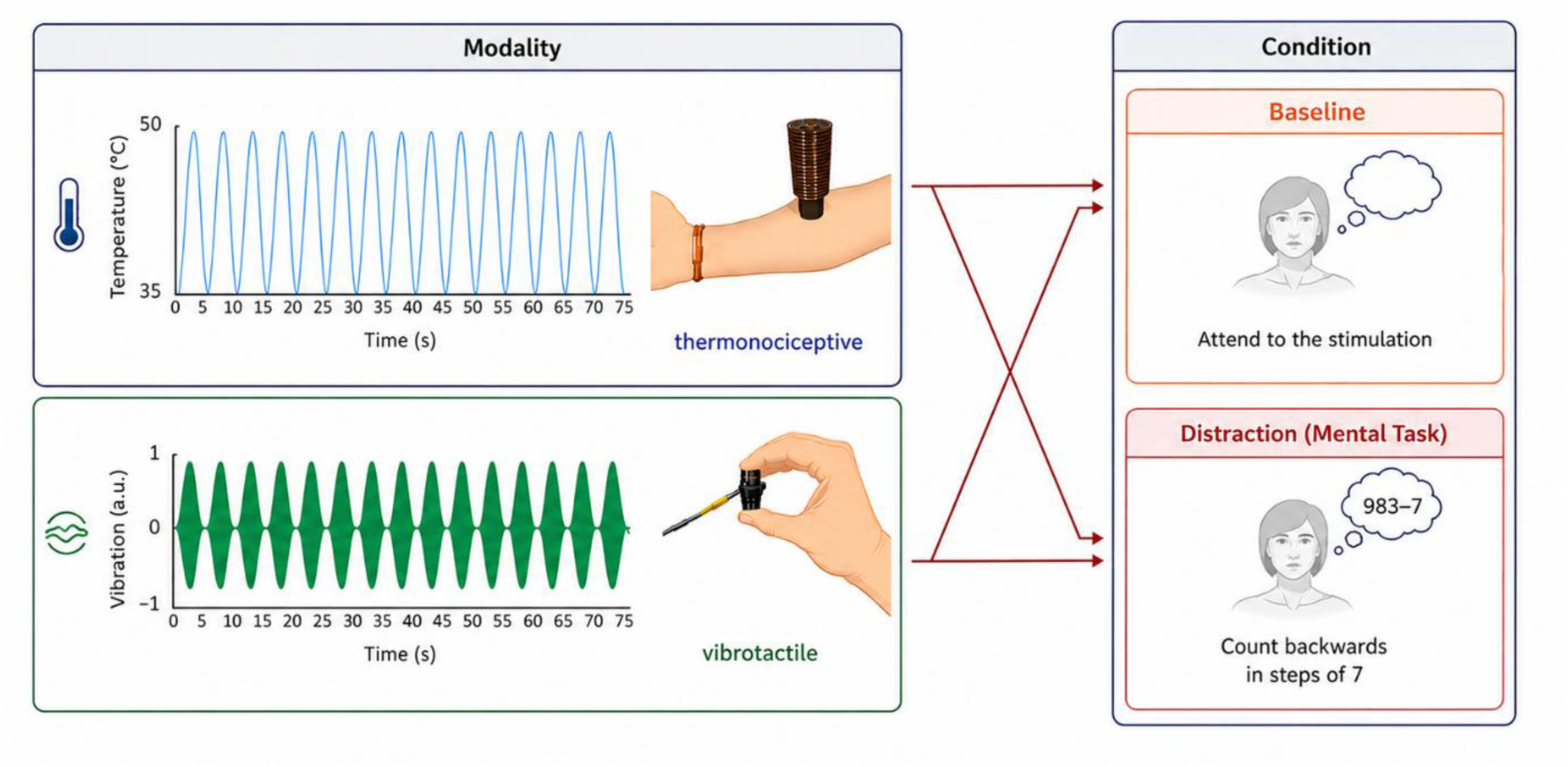
Experimental paradigm, adapted from Leu *et al.* [33]. Each modality (thermonociceptive / vibrotactile stimulation) was paired with each condition (baseline / distraction) for a block of 10 trials. Each trial lasted 75s, during which the stimulation was applied in a sinusoidal manner at 0.2 Hz.

The distraction task consisted of continuously subtracting 7 from a 3-digit starting number throughout the stimulation [19] and to report the final number after the end of the trial. Since our main aim was to keep the patients engaged and motivated for the task, they had the possibility to subtract 3 instead of 7 if they felt like the task was too demanding. This was the case for 2 patients. After each stimulus, the patients provided a rating of the intensity of the stimulus on a numerical ratings scale from 0 to 10 (0: no perception, 10: most intense perception imaginable) and classified the stimulation as painful or not (yes/no).

### 2.3 Electrode implantation

For each patient, the most probable sites of the epileptogenic focus were determined by a multidisciplinary team. Commercially available depth electrodes for stereo electroencephalography (SEEG) (Dixi Medical, Besancon, France) were mostly implanted orthogonally at those sites through burr holes using a frameless neuronavigational approach, as described in Budke *et al.* [8],Joris *et al.* [28]. Post-implantation 3D-1T MRI scans confirmed the accurate placement of each electrode on the day of or the day after the operation.

### 2.4 Electrode contact localization

The exact anatomical location of the implanted electrode rods was determined using BrainVoyager (Brain Innovation, Maastricht, The Netherlands), by normalizing the MRI scans that were acquired post-implantation to a standard T1 template in MNI space and manually identifying the MNI coordinates of the insular electrode contacts in collaboration with the neurology team of the Saint-Luc university hospital. The exact location of the coordinates was then assessed using the Automated Anatomical labelling Atlas 3 [53], running in the open-source toolbox Statistical Parametric Mapping (SPM12) in MATLAB. If there were doubts (i.e. a mismatch between the atlas and the surgical implantation plan), the labelling of the location by the neurology team was preferred. All electrode contacts located in the insula are reported in Table S.1 and are illustrated in Figure 2.

**Figure 2.**
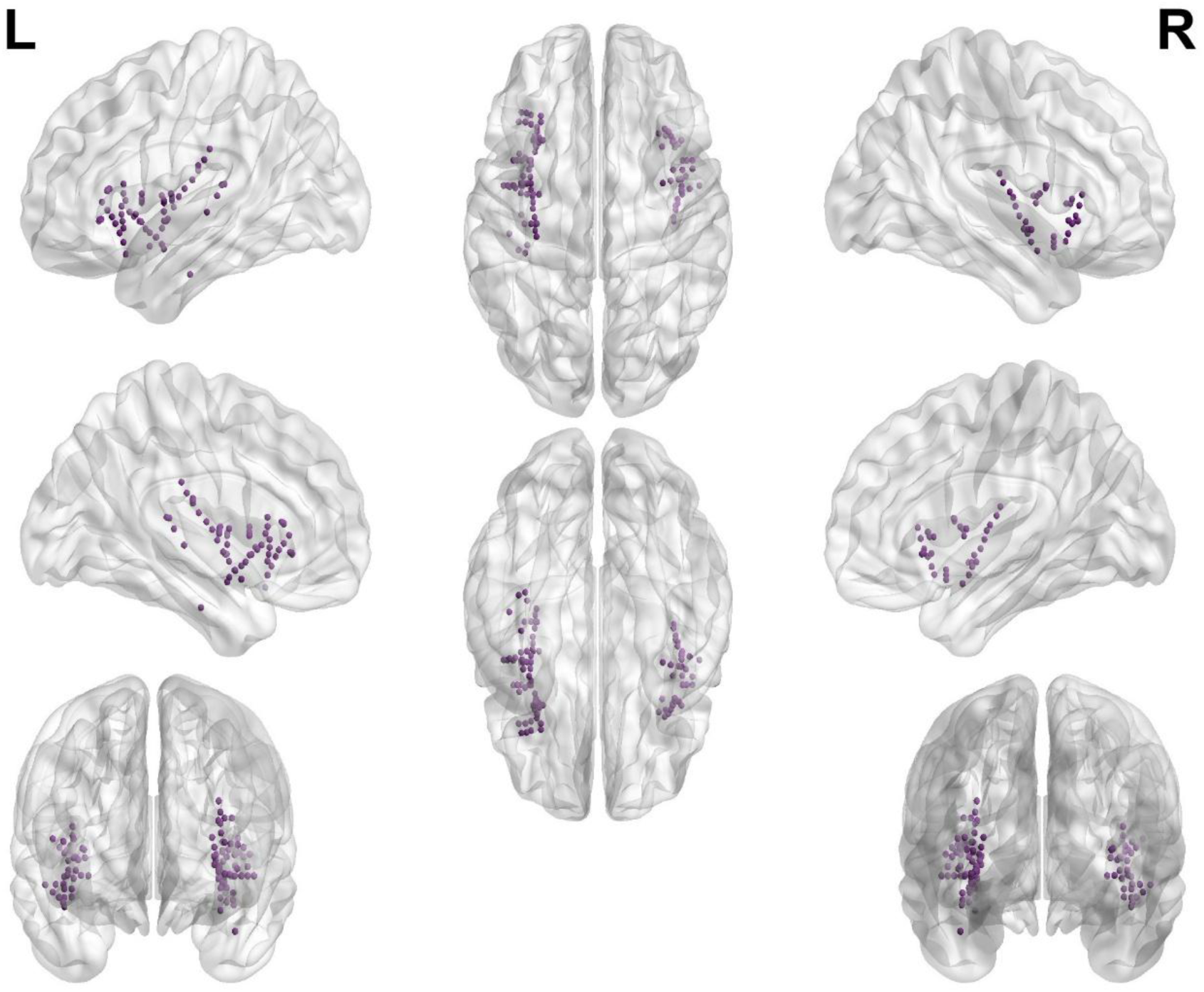
Spatial distribution of intracerebral electrode contacts. Glass-brain representation showing the locations of all electrode contacts included in the analyses.

### 2.5 Intracerebral recordings

In 2 patients, the signal from the implanted electrodes was acquired using a DeltaMed system (Natus Europe, Paris, France). Electromyographic (EMG) and electrocardiographic activity was recorded from the patient’s nondominant arm at the triceps and biceps and from two electrodes respectively located on the right and left side of the sternum, one electrode located centrally under the sternum, and one electrode on the right lateral side, respectively. The reference electrodes were located between Cz and Pz. For all other participants, the signal from the implanted electrodes was acquired using a “BRAIN QUICK ^®^” acquisition system by MicroMed (Natus Europe, Paris, France), due to an update of the systems in place. The reference electrode was placed 2 cm behind the usual Fz location in the 10-20 International System. Additionally, electromyographic (EMG) and electrocardiographic activity (ECG) were recorded to account for artifacts related to movement and heart rate. These bipolar electrodes were placed on each upper limb on the deltoid muscle, as well as in the centre of the chest and on the right upper part of the abdomen. For both systems, the sampling rate was 512 Hz. The two recording setups produced highly comparable data with no apparent difference in data quality. The EEG recordings were analysed offline using the Letswave7 (www.letswave.org) toolbox in MATLAB (2022a The MathWorks).

### 2.6 Preprocessing

A 4^th^ order Butterworth filter over the range of 0.05 to 100 Hz and a notch filter around 50 Hz (width: 2 Hz, to remove line noise) were applied and 75 s epochs were segmented based on the onset of the stimulation. Artifacts related to heartbeat and movements were removed using an Independent Component Analysis (Fast ICA algorithm [27]), leading to the removal of 4 electrodes for each patient, related to EMG and ECG electrodes. The signal was re-referenced to the average of all electrode contacts for each participant separately. The remaining signal was then averaged per participant, condition and modality. To assess the periodic response at the frequency of the stimulation (i.e. “frequency tagging” [51]), the signal was transformed into the frequency domain using a Fast Fourier Transform (FFT) [22], resulting in a resolution of 0.013 Hz in the frequency spectrum. The signal was then baseline-corrected by subtracting the average amplitude measured at the 2-5 surrounding bins, at each electrode and frequency bin and thus largely removing residual noise [43], which is inherently present particularly in lower frequencies. The baseline correction is based on the notion that without the periodic response, the signal at the frequency (bin) of interest should not differ from the average of the surrounding frequency bins.

The modulation of ongoing neural oscillations was assessed in the theta (4-8 Hz), alpha (8-12 Hz), beta (12-40 Hz) and gamma (40-100 Hz) frequency bands using a “frequency-tagging of ongoing oscillations” approach [12,33,34,36,44]. This approach is an extension of the commonly used frequency-tagging [51], applied to ongoing oscillations by filtering the data for the respective frequency band after re-referencing the signal, and estimating the signal envelope using a Hilbert transform. The data was then averaged across trials for each participant and condition/modality separately, before converting the signal into the frequency domain using an FFT. Finally, the signal was baseline corrected as described above. The same preprocessing pipeline has been used by several studies from our lab [12,33,34,36].

### 2.7 Harmonic aggregation

All harmonics in the frequency spectrum were summed up to form the amplitude at the “frequency of interest” (FOI). This procedure is based on the rationale that excluding the harmonics elicited by a periodic stimulation would disregard an important portion of the evoked response [52]. Because the neural response to a periodic stimulus is not perfectly sinusoidal, the elicited activity is distributed over multiple harmonics rather than being centred only at the frequency of stimulation. To this end, harmonics need to be retained to accurately reconstruct the time-domain signal. We therefore summed up all harmonics for each condition and modality, for the phase-locked response as well as for the frequency bands [14,33,34,54]. The amplitudes at these FOIs were consequently used in the statistical analysis.

Additionally, to assess whether potential effects of the distraction task on the modulation of ongoing oscillations were due to differences in magnitude of baseline amplitudes between the modalities, the relative change in the aggregated amplitudes were calculated for each modality and location (Δ amplitude = baseline - distraction).

### 2.8 Statistical analysis

R Statistical Software (V. 4.3.1 [49]) was used for the statistical analysis of the data. A Holm-corrected right-tailed Wilcoxon signed-rank test was performed at the FOI for the phase-locked response as well as the modulation of ongoing oscillations in the frequency bands, to assess whether a statistically significant response (i.e. amplitude larger than zero) was present.

Separate linear mixed-effects models (LMM) were fitted for each outcome variable. The model for perceived stimulation intensity included condition, modality, and their interaction as fixed effects. Models for the EEG outcomes included condition, modality, electrode location (anterior/posterior), and their two-and three-way interactions as fixed effects and were fitted separately for the periodic EEG response and the modulation of ongoing oscillatory activity within the frequency bands of interest. All models were fitted using restricted maximum likelihood (REML). Random intercepts were included for patients in the intensity analyses and for electrode-within-patient combinations in the EEG analyses to account for the hierarchical structure of the data and clustering of observations within patients and electrodes. Significant results (p<0.05) were further assessed using post-hoc pairwise comparisons of the model-estimated marginal means, Tukey-adjusted p-values are reported for multiple comparisons.

All EEG amplitudes were transformed using the Yeo-Johnson transformation [69] to conform to the assumptions underlying the LMMs. This specific transformation method was chosen as it can also handle negative values, which is not the case for the log-transformation. After transformation, influential data points were objectively identified using Cook’s distance [D] [13], and removed if D>1. Given these guidelines, only few outliers (i.e. single datapoints) were removed from the analysis (PL, beta, gamma: 0, theta: 1, alpha: 2).

To assess whether the absence of a condition × modality × orientation interaction reflected a true lack of effect or insufficient sensitivity, a Bayesian linear mixed model was used to quantify the posterior distribution of the interaction term. A region of practical equivalence was defined a priori as β = [−0.1, 0.1] on the log scale (∼ ±10% change), corresponding to a negligible effect size in the outcome metric.

The relative (Δ = baseline - distraction) amplitude between the conditions was tested against 0 using a Wilcoxon signed-rank test (Holm-corrected) to assess whether a difference between the conditions was present regardless of baseline amplitude magnitude.

As an additional exploratory analysis, we used LMM’s (relAmplitude∼axis*modality+(1|patient)) to investigate whether gradients along the X (left/right), Y (anterior/posterior) and Z axis (dorsal/ventral) of the MNI coordinates electrode contact positions could explain the relative difference between baseline and distraction amplitudes and support the potential findings of the previous analysis where anterior / posterior was assigned. One model was run for each axis and each frequency band.

## 3. Results

### 3.1 Task performance and perception

The reported number after the arithmetic task was correct in 51.4% (36/70) during vibrotactile and in 36.1% (35/97) of answers during thermonociceptive stimulation. 2 participants did not complete the vibrotactile part of the study, and the task data of one patient has been corrupted by a technical issue. A paired one-tailed t-test showed that the task performance did not differ between modalities (t(6)=-1.0, p=0.178, d=0.17). On average, participants subtracted 43.4 ± 45.5 (mean ± std dv) steps during thermonociceptive stimulation, and 54.9 ± 58.4 steps during vibrotactile stimulation (t(6)=-0.591, p=0.288, d=0.002).

Ratings varied significantly between modalities (F(1,336.35)=42.662, p<0.001, η^2^ =0.113)) and conditions (F(1,335)=25.256, p<0.001, η^2^ =0.070) (Figure 3A). The interaction between modality and condition was not significant (F(1,335)=0.007, p=0.933, η^2^ =0.000). The thermonociceptive stimuli were perceived as painful during less than half of the baseline condition (41%), and to an even lesser extent during distraction (23%). Vibrotactile stimuli were never perceived as painful in either condition (0%). Comparing the conditions in the thermonociceptive modality (Figure 3B) revealed a significant difference between the number of stimuli perceived as painful during baseline and distraction (W(10)=28, p=0.022, r=0.806).

**Figure 3.**
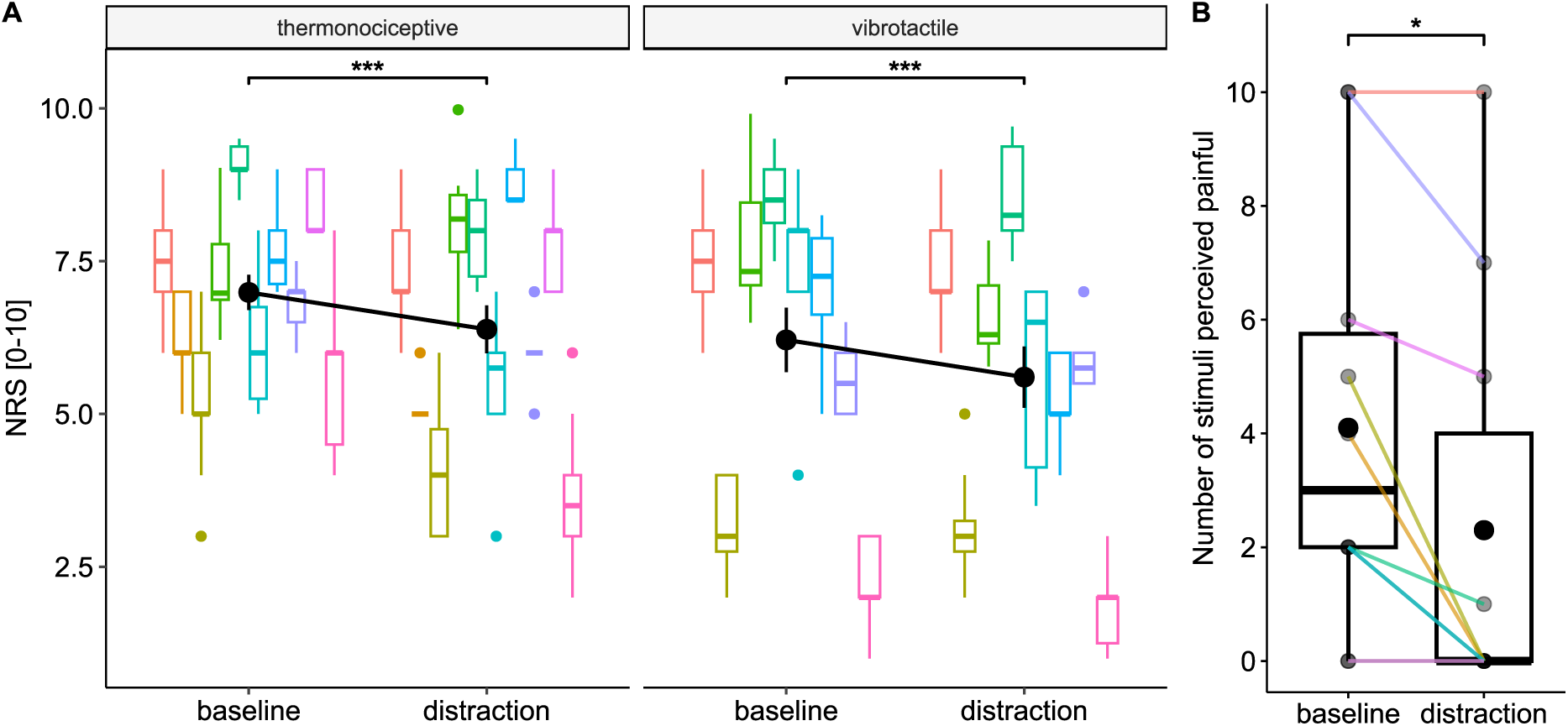
Perception ratings. A: Perceived stimulus intensity using a Numerical Rating Scale (NRS) between 0 (“no perception”) and 10 (“most intense perception imaginable”) for thermonociceptive and vibrotactile stimulation. Coloured horizontal lines mark the individual average, black lines indicate group means. Significant main effects of modality and condition were observed in the Linear Mixed Model (***p<0.001). B: Number of stimuli that were rated as painful per participant and condition in the thermonociceptive modality. The p-value reflects Wilcoxon paired signed-rank test (*p<0.05).

### 3.2 Neural response

#### 3.2.1 Wilcoxon test

A significant periodic phase-locked response was found at the FOI for both modalities, conditions and insular regions, except for the anterior thermonociceptive distraction response (Supplementary Table S.2). Figure 4 illustrates the neural response in the first 1 Hz of the frequency spectrum for thermonociceptive and vibrotactile stimulation in the anterior and posterior insula. Similarly, the modulation of ongoing oscillations at the FOI was significantly different from zero in the theta and alpha frequency bands for all tested factors apart from the amplitude at the FOI exerted by thermonociceptive stimulation in the anterior insula. In the beta frequency band, significant modulation of ongoing oscillations was observed predominantly in the posterior insula, with the addition of the vibrotactile response during distraction in the anterior insula. Finally, in the gamma frequency band, the thermonociceptive stimulation led to significant responses at the FOI in all tested factors except for in the anterior insula during distraction. For vibrotactile stimuli, significant responses were found in the anterior insula during distraction as well as in the posterior insula during baseline.

**Figure 4.**
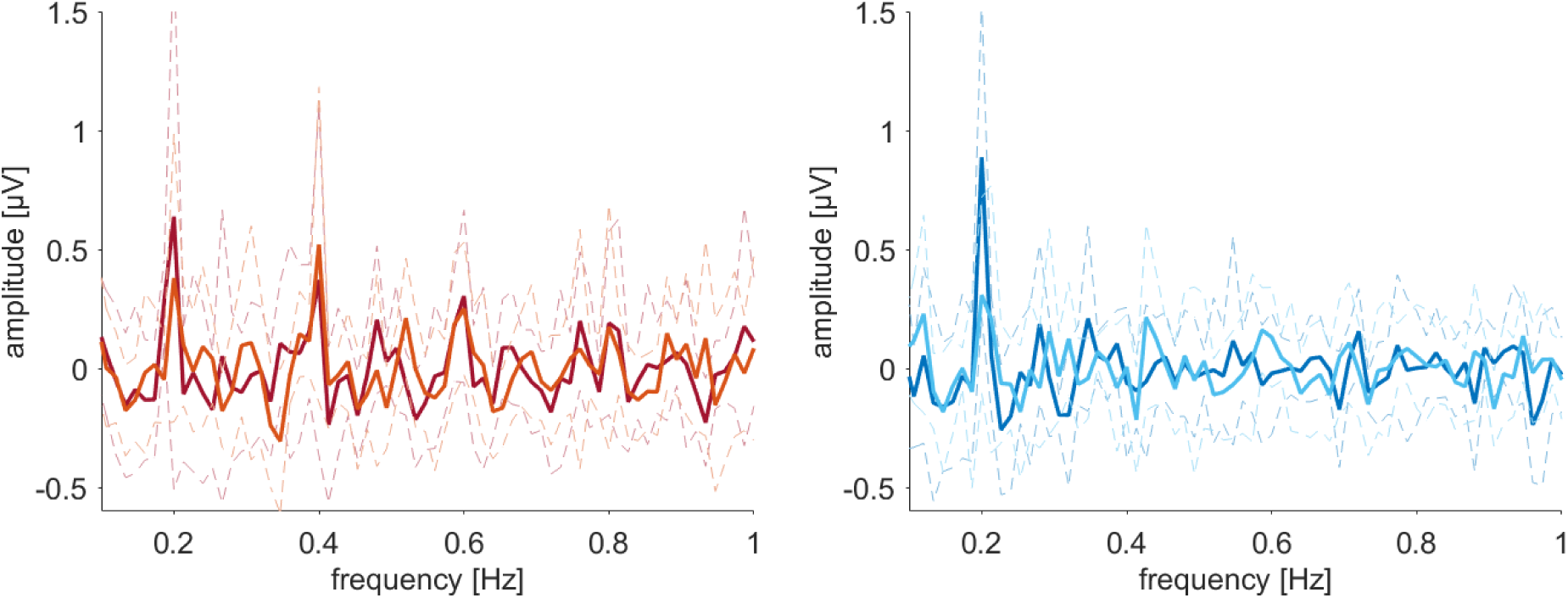
Group average of the phase-locked response across the frequency spectrum elicited by slow sustained periodic thermonociceptive (A) and vibrotactile (B) stimulation at 0.2 Hz and its first few harmonics in the anterior insula. The lighter shaded line indicates the distraction condition, while dotted lines mark the standard deviation from the mean.

#### 3.2.2 Linear mixed models

The complete results of the LMMs are summarized in Table 1. Similar results were obtained across all frequency bands and the phase-locked response; the main effect of *location* was significant, as was the interaction between *location* and *modality* (except for the beta frequency band), indicating that the location of the electrode contact (anterior vs posterior) significantly influenced the elicited amplitudes. Post-hoc pairwise comparisons of estimated marginal means (detailed comparison in supplementary table S.3, illustrated in figures 5A for the phase-locked response and Figure 6 for the modulation of ongoing oscillations) demonstrated that neural responses elicited by thermonociceptive stimulation were systematically larger than vibrotactile responses in the posterior insula. Interestingly, in the anterior insula, in the theta and alpha frequency band, the opposite relationship was found, with vibrotactile responses being significantly larger than thermonociceptive ones. Further, significant main effects for *condition* were found in both theta and beta frequency band, showing a larger modulation in the distraction condition. A significant interaction between *condition* x *modality* was only found in the beta frequency band (table 1), and post-hoc pairwise comparison revealed a significantly larger response during distraction compared to baseline following vibrotactile stimulation (EMM=0.11, t(241.23)=-2.83, p=0.01, d=-0.36). No such difference was found in the thermonociceptive modality (EMM=-0.01, t(241.23)=-0.23, p=0.82, d=-0.03).

**Figure 5.**
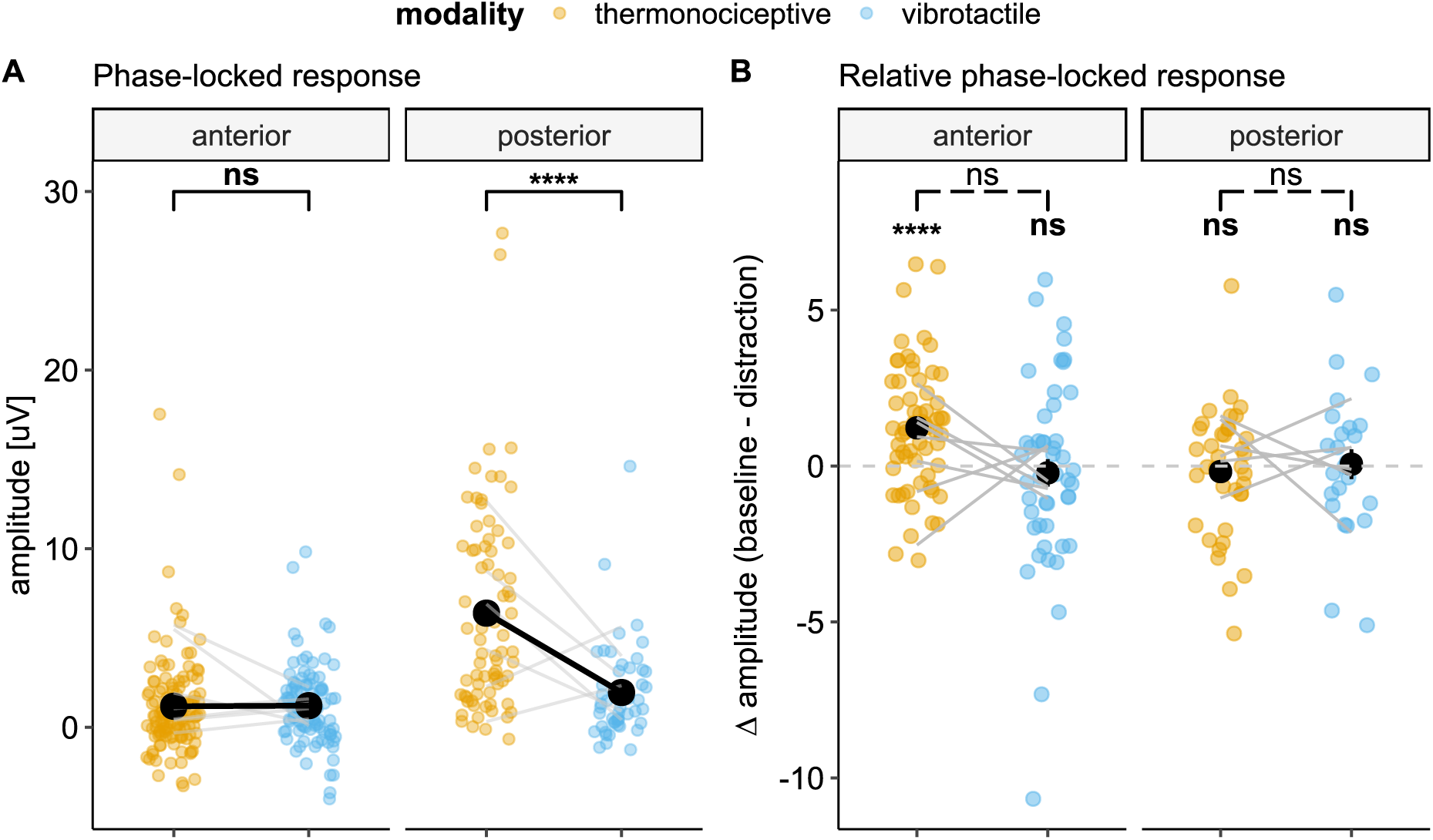
Phase-locked response. Panel A) illustrates the post-hoc pairwise comparison (Tukey-adjusted) of the interaction between condition and modality on the aggregated EEG amplitudes. Panel B) represents the relative amplitude (distraction subtracted from the baseline amplitude). Individual data are represented in coloured dots, while group means are represented in black. Grey horizontal lines mark the average for each patient. *p<0.05, **p<0.01, ***p<0.001, ****p<0.0001.

**Figure 6.**
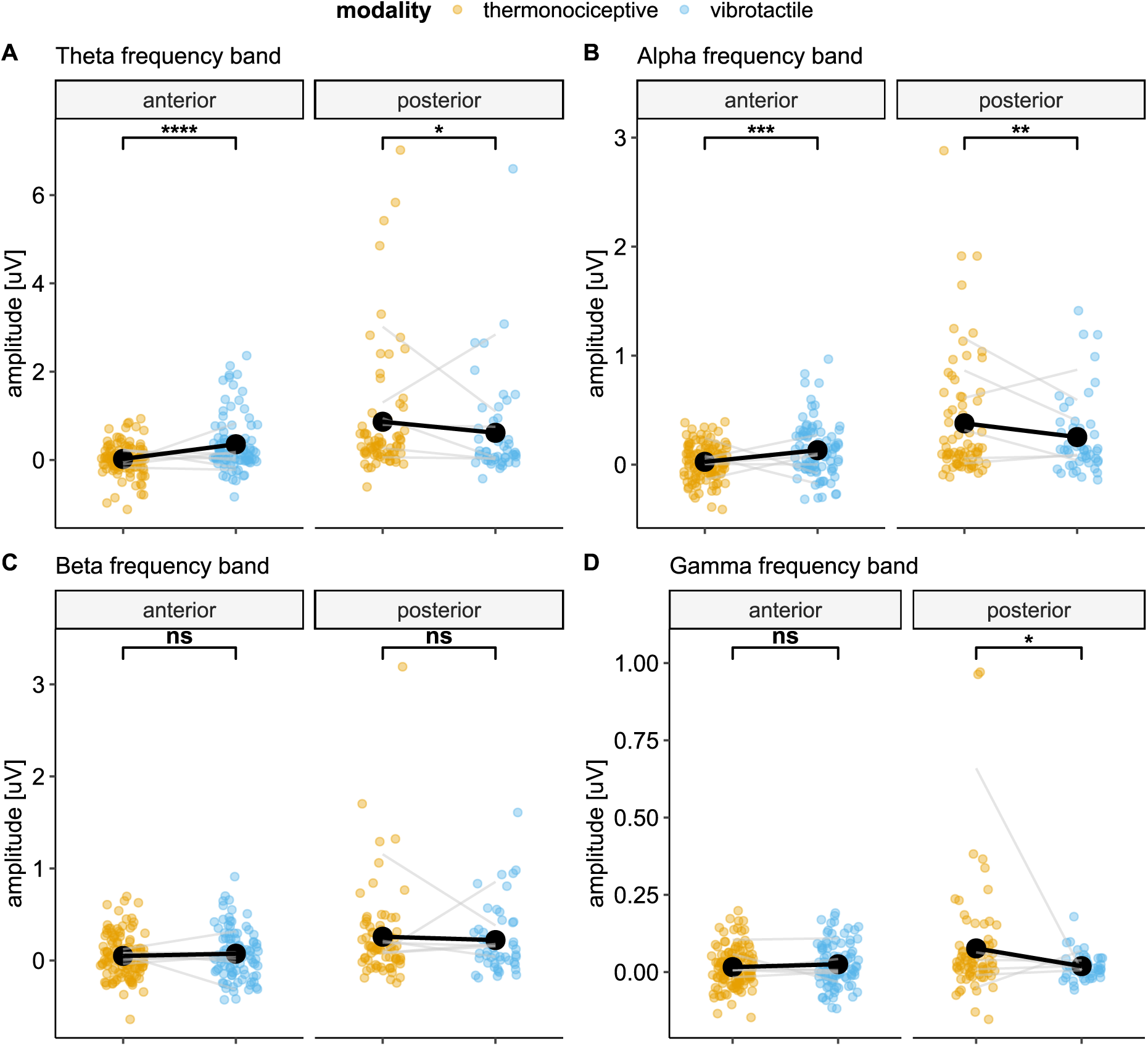
Post-hoc pairwise comparison of the LMM interaction between condition and modality (Tukey-adjusted). Individual data are represented in coloured dots, while group means are represented in black. Grey horizontal lines mark the average for each patient. *p<0.05, **p<0.01, ***p<0.001, ****p<0.0001.

**Table 1.** Results of the Linear Mixed Model (amplitude ∼ modality * condition * location + (1| electrode contact:patient) and estimated partial effect sizes. Significant results (p<0.05) are indicated in bold font.

|  | *Parameter* | *df* | *F* | *p* | *eta*<sup>2</sup><sub>*p*</sub> |
| --- | --- | --- | --- | --- | --- |
| *Phase-locked response* | condition | 1, 240.424 | 1.203 | 0.274 | 0.005 |
|  | **modality** | **1, 279.435** | **25.127** | **0.000** | **0.083** |
|  | **location** | **1, 99.429** | **38.955** | **0.000** | **0.282** |
|  | condition:modality | 1, 240.424 | 0.812 | 0.368 | 0.003 |
|  | condition:location | 1, 240.424 | 2.831 | 0.094 | 0.012 |
|  | **modality:location** | **1, 279.435** | **25.188** | **0.000** | **0.083** |
|  | condition:modality:location | 1, 240.424 | 0.327 | 0.568 | 0.001 |
| *Theta* | **condition** | **1, 238.805** | **5.146** | **0.024** | **0.021** |
|  | modality | 1, 273.654 | 1.149 | 0.285 | 0.004 |
|  | **location** | **1, 100.159** | **16.472** | **0.000** | **0.141** |
|  | condition:modality | 1, 238.805 | 1.796 | 0.181 | 0.007 |
|  | condition:location | 1, 238.805 | 1.814 | 0.179 | 0.008 |
|  | **modality:location** | **1, 273.654** | **22.709** | **0.000** | **0.077** |
|  | condition:modality:location | 1, 238.805 | 0.000 | 0.991 | 0.000 |
| *Alpha* | condition | 1, 237.484 | 2.981 | 0.086 | 0.012 |
|  | modality | 1, 264.777 | 0.105 | 0.747 | 0.000 |
|  | **location** | **1, 101.348** | **19.161** | **0.000** | **0.159** |
|  | *condition:modality* | *1, 237.419* | *3.474* | *0.064* | *0.014* |
|  | *condition:location* | *1, 237.484* | *3.528* | *0.062* | *0.015* |
|  | **modality:location** | **1, 264.777** | **19.592** | **0.000** | **0.069** |
|  | condition:modality:location | 1, 237.419 | 1.337 | 0.249 | 0.006 |
| *Beta* | **condition** | **1, 241.234** | **5.391** | **0.021** | **0.022** |
|  | modality | 1, 283.684 | 0.011 | 0.918 | 0.000 |
|  | **location** | **1, 98.283** | **16.235** | **0.000** | **0.142** |
|  | **condition:modality** | **1, 241.234** | **4.134** | **0.043** | **0.017** |
|  | condition:location | 1, 241.234 | 0.019 | 0.890 | 0.000 |
|  | modality:location | 1, 283.684 | 0.426 | 0.514 | 0.002 |
|  | condition:modality:location | 1, 241.234 | 0.002 | 0.964 | 0.000 |
| *Gamma* | condition | 1, 239.943 | 0.726 | 0.395 | 0.003 |
|  | modality | 1, 278.331 | 1.972 | 0.161 | 0.007 |
|  | **location** | **1, 100.477** | **4.116** | **0.045** | **0.039** |
|  | condition:modality | 1, 239.943 | 2.663 | 0.104 | 0.011 |
|  | condition:location | 1, 239.943 | 0.001 | 0.981 | 0.000 |
|  | **modality:location** | **1, 278.331** | **6.012** | **0.015** | **0.021** |
|  | condition:modality:location | 1, 239.943 | 2.048 | 0.154 | 0.008 |

Given the absence of a triple-interaction effect between condition, modality, and location, post-hoc Bayesian inference was used to assess whether there is evidence for absence of this interaction effect. Across all analyses, the condition × modality × location interaction was consistently centred near zero. In the phase-locked response, the interaction was highly uncertain (β = −0.41, 95% CrI [−1.80, 0.97], P(|β| < 0.1) = 0.092), indicating insufficient precision to distinguish small effects from true absence. In the theta band, estimates were similarly centred near zero (β = 0.00, 95% CrI [−0.33, 0.33], P(|β| < 0.1) = 0.48), again providing weak evidence for practical equivalence. In contrast, the oscillatory bands showed more precise and consistently small effects. In the alpha and beta bands, estimates were close to zero (alpha: β = −0.09, 95% CrI [−0.25, 0.07], P(|β| < 0.1) = 0.49; beta: β = 0.01, 95% CrI [−0.20, 0.22], P(|β| < 0.1) = 0.83), indicating moderate support for practical equivalence. The strongest evidence for absence was observed in the gamma band, where the interaction was tightly constrained around zero (β = −0.04, 95% CrI [−0.10, 0.02]) with a high probability of lying within the equivalence region (P(|β| < 0.1) = 0.973). Overall, these results converge with the frequentist findings, providing no evidence for a meaningful higher-order interaction and suggesting that any potential effects are small in magnitude.

#### 3.2.3 Exploratory analysis I: Relative change in amplitude

Finally, the comparison of the relative change in amplitude (baseline - distraction) between the conditions showed that in the posterior insula, the distraction task never led to a significant change in relative amplitude in either of the applied stimulation modalities (Figure 7, detailed comparison in supplementary table S.4). However, in the anterior insula, a significant positive increase in the relative amplitude indicates that the distraction task led to a reduction in neural response at the FOI in the phase-locked response (figure 4B). Conversely, the distraction task led to significantly larger neural responses (i.e. negative relative amplitude) in the theta, alpha and gamma frequency bands. No difference was observed between the modalities in the posterior insula (*phase-locked*: t(23)=0.666, p=0.512, d=0.136); *theta:* t(23)=1.439, p=0.163, d=0.294*; alpha:* t(21)=1.390, p=0.179,d=0.296*; beta:* t(23)=2.088, p=0.096, d=0.426; *gamma:* t(21)=0.922, p=0.367, d=0.197).

**Figure 7:**
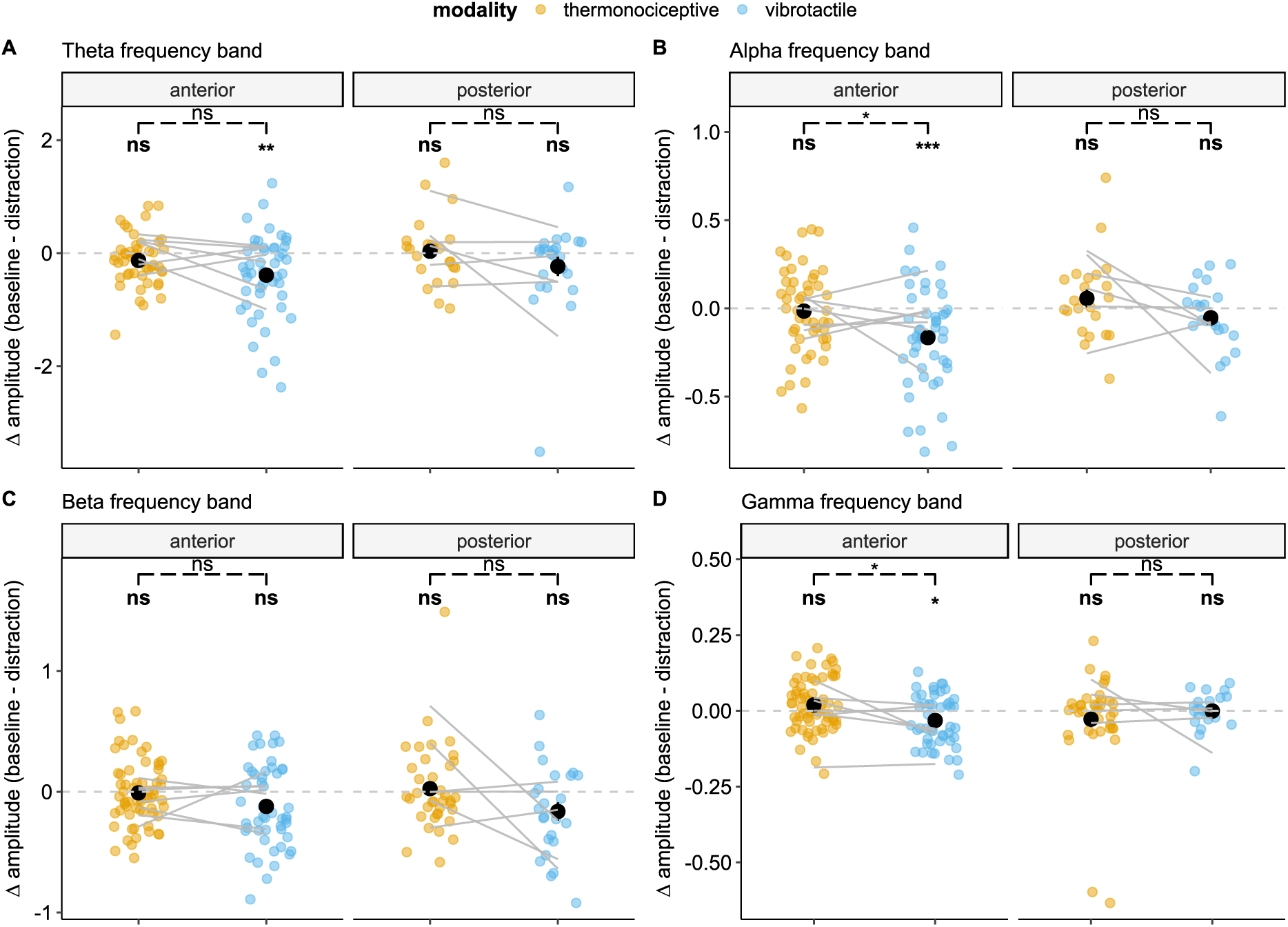
Relative (Δ) amplitudes (baseline - distraction) were computed to evaluate potential baseline influences on effects identified by the LMM. Group means and confidence intervals are shown in black. One-sample t-tests (Holm-corrected) assessed whether relative amplitudes differed zero (**^ns^**p>0.05, **\***p<0.05, **\*\***p<0.01, **\*\*\***p<0.001), thereby testing the presence of a distraction effect independent of amplitude magnitude. Pairwise t-tests (Holm-corrected) were conducted to compare relative amplitudes between modalities (^ns^p>0.05, *p<0.05).

In the anterior insula, a difference between modalities was found in the alpha and gamma frequency band, although directed by a negative relative amplitude (i.e. increase in neural response during distraction) in the vibrotactile modality (*alpha*: t(47)=2.599, p=0.024, d=0.375; *gamma*: t(47)=2.854, p=0.012, d=0.412). No such difference was found between the modalities in the phase-locked response ((t(47)=2.274, p=0.055, d=0.328), beta (t(47)=0.830, p=0.411, d=0.120) and theta (t(46)=1.923, p=0.061, d=0.0.281) frequency band after correction for multiple comparisons.

#### 3.2.4 Exploratory analysis II: Relative amplitude gradients along insular axes

In this exploratory analysis, we examined if the modality effects identified in the LMMs varied along the anatomical axes of the insula by testing whether interactions between MNI electrode coordinates along the x-, y-or z-axis and modality could explain the variance in the relative EEG amplitudes. In the alpha frequency band, a significant interaction between modality and anterior-posterior (y-axis) coordinates was found (F(1, 158.24)= 6.90, p=0.01, η²p=0.040). Post-hoc pairwise comparisons revealed a significant difference between modalities along the anterior–posterior axis (Figure 8; EMM=0.006, t(159)=2.626, p= 0.010, d=0.42). A similar pattern was observed in the beta frequency band along the ventral-dorsal axis, with a significant interaction between modality and axis coordinates (F(1159,78)=4.162, p=0.043, η²p=XX, η²p=0.030). Relative amplitudes elicited by vibrotactile stimulation varied significantly along the ventral–dorsal axis, whereas thermonociceptive responses did not (Figure 8; EMM=0.010, t(160)=2.040, p=0.043, d=0.32).

**Figure 8.**
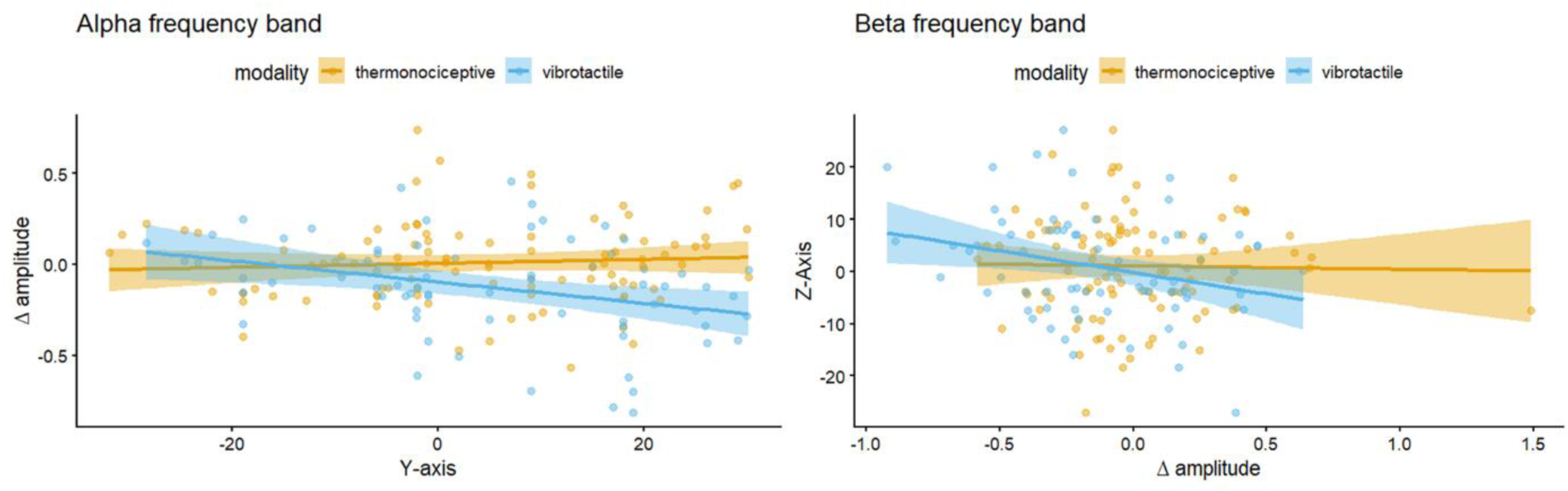
Gradient analysis. Illustrating the relationship between A) the relative amplitude measured in the alpha frequency band and the electrode contact location along the anterior-posterior axis and B) the relative amplitude measured in the beta frequency band and the electrode contact locations along the dorsal-ventral axis.

## 4. Discussion

The present study tested whether distraction-related reductions in perceived stimulus intensity are mirrored by changes in frequency-tagged insular responses to sustained thermonociceptive and vibrotactile stimulation. Although the arithmetic distraction task reduced perceived stimulus intensity in both modalities, this perceptual modulation was not accompanied by a reduction of oscillatory responses in the posterior insula. Thus, contrary to our initial hypothesis, responses in the posterior insula captured by frequency tagging did not track changes in subjective intensity induced by distraction.

A periodic modulation of ongoing oscillations was found across both stimulation modalities in the anterior and posterior insula, with consistently larger modulation in the posterior insula. These findings largely align with Liberati *et al.* [36], who reported preferential modulation of ongoing alpha-and theta-band oscillations in the dorsal-posterior insula following thermonociceptive stimulation. However, we also observed significant periodic responses following vibrotactile stimulation. This discrepancy may reflect methodological differences, as we aggregated harmonics across the frequency spectrum to quantify the full neural response [52]. Differences in stimulus quality may also have contributed; whereas the laser stimuli used by Liberati *et al.* [36] were generally perceived as painful, the thermonociceptive stimuli in the present study were mostly perceived as non-painful, potentially reflecting differences in the efficacy of nociceptive activation between laser and contact-heat stimulation.

At the level of scalp EEG, a previous study using the identical experimental paradigm did not show condition-related neural modulation effects despite a comparable decrease in perceived stimulation intensity [33]. This likely reflects the predominant sensitivity of scalp EEG to somatosensory cortices [2,19] and a resulting insensitivity to signals originating from deeper structures such as the insula [50]. In contrast, intracerebral recordings in the present study reveal clear anterior insula modulation effects, suggesting that distraction-related changes in somatosensory processing are not fully captured at the scalp level.

Consistent with Liberati *et al.* [36] and the higher perceived intensity of thermonociceptive stimulation at baseline, thermonociceptive responses in the posterior insula were systematically larger than vibrotactile responses. Because this preferential modulation was observed even when thermonociceptive stimuli were largely non-painful, the findings suggest that the modulation of ongoing oscillations in the posterior insula is not specific to pain perception, but rather more closely related to stimulus intensity. This interpretation is further supported by the task effects on the modulation of ongoing oscillations: no reduction in thermonociceptive modulation was observed during the arithmetic task, despite a decrease in perceived intensity. Although larger baseline responses may have had greater capacity for change, analyses of relative differences between baseline and distraction conditions yielded comparable results. Modality differences observed in the alpha and gamma bands were driven by a relative increase in vibrotactile modulation in the anterior insula during the arithmetic task, whereas no corresponding change was observed for thermonociceptive stimulation. Together, these findings suggest that modulation of ongoing oscillations primarily reflects objective stimulus properties, such as intensity, rather than subjective perception or pain-specific processing.

Previous evidence suggests a predominantly posterior involvement of the insula in the processing of nociceptive stimulation [31]. Several investigations studied the effect of cognitive distraction on pain perception in relation to insular activity and generally found a decrease during distraction from nociceptive stimulation concomitant to a decrease in pain perception [5,47,48,58,65]. No such decrease in posterior insula activity was observed despite reduced perceived intensity during distraction, suggesting a closer relationship between posterior insular oscillations and stimulus intensity than stimulus perception. However, this interpretation should consider the frequency-tagging approach, which enhances signal-to-noise ratio for strictly periodic responses but necessarily reduces sensitivity to non-periodic or temporally evolving effects. In the present design, where the distraction task extended over 75 seconds with a single post-stimulus rating, any dynamic changes in cognitive state over time cannot be resolved. As a result, frequency-tagging may not capture transient or non-repetitive aspects of the distraction effect on ongoing oscillatory activity. However, this explanation does not readily account for the presence of modulation in the vibrotactile modality. Accordingly, modality-specific differences in salience or network engagement may also contribute to the observed dissociation.

The anterior and posterior insula differ in both structural and resting state connectivity with other brain regions involved in pain perception, with pain awareness being related mostly to the anterior insula [67]. Similarly, cognitive factors influencing pain perception such as attention and expectation have been shown to be integrated with the objective properties of the nociceptive stimulation in the anterior insula [3]. Other investigations showed that, while nociceptive stimulation elicited activity in the posterior part of the insula, distraction mostly attenuated activity in the anterior insula [7,60]. Our results support this view, as distraction-related effects following thermonociceptive stimulation were restricted to phase-locked responses in the anterior insula, without corresponding changes in modulation of ongoing oscillations. Because phase-locked activity reflects stimulus-evoked responses with consistent timing relative to stimulus onset, this suggests that cognitive distraction primarily reduced the precision or synchrony of thermonociceptive encoding, likely reflecting decreased allocation of attentional resources to salient inputs in the anterior insula. In contrast, modulation of ongoing oscillations elicited by thermonociceptive stimulation appears to index more distributed and variable network dynamics that were not significantly affected by the present manipulation. If the observed dissociation between neural modulation and perceived intensity reflects a general mechanism rather than a thermonociception-specific effect, a similar pattern should be evident for vibrotactile stimulation.

Without any task, vibrotactile stimulation was shown to primarily activate the primary and secondary somatosensory cortices, as well as the posterior insula [1,20,46]. Yet, if a working memory task such as a frequency discrimination task is being performed, fMRI investigations have demonstrated that activity shifts to the anterior insula [1,35,62]. Considering that the insula receives input from somatosensory cortices [4], which are then relayed along the posterior-anterior axis of the insula to higher-order cortical structures such as the prefrontal region [10,64], the observed effects are likely related to cognitive task monitoring and short-term memory retention of the perceived tactile stimulation which are integrated in the anterior insula [9]. Additionally, studies examining the circuits involved in emotional regulation found the anterior insula to be active during distraction tasks, e.g. mental arithmetic tasks [18,41]. The observed increase in modulation during vibrotactile stimulation in the anterior insula matches these reports, pointing towards an increase in activity due to cognitive effort.

But why were these changes not observed during thermonociceptive stimulation? One potential explanation is a mismatch in saliency between the modalities. Painful stimuli are inherently salient due to their potential threat value, and at least a subset of the thermonociceptive stimuli were perceived as such, increasing salience compared to vibrotactile stimulation. As a core node of the salience network [40,68], the anterior insula may therefore have been more strongly engaged by the thermonociceptive input, potentially interfering with resource allocation to the arithmetic task. While task performance did not differ between modalities, subjective reports indicated that thermonociceptive stimulation intermittently disrupted focus and calculation rhythm, whereas vibrotactile stimulation was less intrusive.

To further explore anterior-posterior differences and recent evidence suggesting a graded organization of the insula [55,66], we examined whether anterior-posterior, ventral-dorsal or left-right gradients explain modality differences in relative EEG amplitude as an exploratory analysis. The presence of spatial gradients for vibrotactile responses may help explain why cognitive manipulation through distraction preferentially affected this modality. If vibrotactile processing engages a broader range of insular regions, including anterior contacts that are more sensitive to task demands, modulation by distraction may become more readily detectable. In contrast, no comparable spatial gradient was observed for thermonociceptive stimulation. This result mirrors the absence of significant modulation of thermonociceptive EEG responses during distraction, despite reduced intensity ratings. Together, these findings suggest a relative stability of thermonociceptive frequency-tagged responses across the sampled insular contacts and experimental conditions. Rather than being fully captured by categorical anterior, middle, and posterior subdivisions, modality effects may vary continuously across insular space. However, given the exploratory nature of these analyses and the heterogeneous spatial sampling inherent to intracranial recordings, these observations should be interpreted cautiously and require replication in larger datasets.

The intracerebral recordings provide valuable data but introduce several limitations. First, electrode coverage within the insula was unbalanced, with more contacts in the anterior than posterior region, and variability in the number of contacts per patient, which may have biased results toward individuals with higher coverage. However, excluding the two patients with the highest number of contacts did not substantially alter the main findings. Additionally, task performance metrics may not accurately reflect participants’ level of distraction, as incorrect responses do not necessarily reflect disengagement or failure of task execution.

## 5. Conclusion

Cognitive distraction reduced perceived stimulation intensity across thermonociceptive and vibrotactile modalities, but contrary to our hypothesis, this behavioural effect was not mirrored by posterior insula activity. Instead, insular oscillatory responses were region-and modality-specific, revealing a dissociation between anterior and posterior insula during distraction, with posterior activity primarily tracking stimulus intensity and anterior responses showing distraction-related modulation. These findings highlight distinct insular contributions to thermonociceptive and vibrotactile processing and underscore the value of intracranial EEG frequency-tagging for disentangling their neural dynamics.

## Acknowledgements

We wish to thank Julien Lambert of the CATL/NTMD team of the Université catholique de Louvain for technical assistance. We would further like to thank all the master’s students that were interns of the lab between 2022-2025 and helped with data acquisition.

During the preparation of this article, the authors used a Large Language Model (“ChatGPT”) to improve grammar and clarity of the language. After using this tool, the authors reviewed and edited the content as needed and take full responsibility for the content of the publication.

## Author contributions (CRediT)

- Conceptualization: CL, GL
- Methodology (experiment): CL, GL
- Methodology (surgical implantation of intracerebral electrodes): VJ, PF
- Methodology (localization of intracerebral electrodes): AB, CL
- Investigation: CL, AB
- Formal analysis: CL
- Visualization: CL, AB
- Funding acquisition: GL, CL
- Resources (patients): SFS, AF
- Supervision: GL
- Writing – original draft: CL
- Writing – review and editing: CL, AB, SFS, AF, VJ, PF, GL

