## Supplementary Materials for "Posterior insula oscillations are not modulated by changes in perceived intensity: Evidence from human intracerebral EEG"

### Supplementary Materials
### for

#### Posterior insula oscillations are not modulated by changes in perceived intensity: Evidence from human intracerebral EEG

Chiara Leu[1,2*], Aïcha Boutachkourt [1,3], Susana Ferrao Santos[1,4], Alexane Fierain[4], Vincent Joris[1,5], Patrice Finet[5], Giulia Liberati[1,6]

[1] Institute of Neuroscience, Université catholique de Louvain, Brussels, Belgium

[2] Center for Interdisciplinary Pain Medicine, Department of Neurology and TUM-Neuroimaging Center, School of Medicine and Health, Technical University of Munich, Munich, Germany

[3] The Leuven Brain Institute, KU Leuven, Leuven, Belgium

[4] Department of Neurology, Saint-Luc University Hospital, Brussels, Belgium

[5] Department of Neurosurgery, Saint-Luc University Hospital, Brussels, Belgium

[6] Psychological Sciences Research Institute, Université catholique de Louvain, Louvain-la-Neuve, Belgium

*Corresponding author: Chiara Leu, Center for Interdisciplinary Pain Medicine, Technical University of Munich, Ismaninger Str. 22, Munich 81675, Germany.

| patient | side | electrode | contact | location | MNI -X | MNI-Y | MNI-Z |
|---|---|---|---|---|---|---|---|
| **p2** | left | INA | 1 | anterior | 31 | 21 | -3.88 |
| | | | 2 | anterior | 34.44 | 20.06 | -3.69 |
| **p3** | right | GTS_1 | 1 | posterior | 42.8 | -5.9 | -7.3 |
| | | | 2 | posterior | 46.57 | -3.57 | -7.48 |
| | | | 3 | posterior | 51.6 | -2.13 | -7.6 |
| | | INA_1 | 1 | anterior | 41 | 16.27 | -3.64 |
| **p4** | right | LOP | 1 | posterior | -39 | -2 | 0.2 |
| | | | 2 | posterior | -43 | -2 | 4 |
| | | | 3 | posterior | -46 | -2 | 5 |
| | | LIA | 1 | anterior | -38.29 | -2 | -16 |
| | | | 2 | anterior | -38 | -1 | -14 |
| | | | 3 | anterior | -38 | 2 | -11 |
| | | | 4 | anterior | -35 | 5 | -9 |
| | | | 5 | anterior | -34 | 7.11 | -7.33 |
| | | | 6 | anterior | -34 | 9.11 | -4.33 |
| | | | 7 | anterior | -32 | 12.88 | -1.88 |
| | | | 8 | anterior | -31.5 | 15 | 0 |
| | | LOF | 1 | anterior | 33 | -1 | 7 |
| | | | 2 | anterior | -36 | -1.13 | 7.13 |
| | | | 3 | anterior | -39 | -1 | 7.14 |
| | | LT | 1 | anterior | -42 | -3.13 | -3.88 |
| | | | 2 | anterior | -45 | -2.67 | -3.89 |
| | | | 3 | anterior | -48 | -2.13 | -3.88 |
| **p5** | right | H | 1 | posterior | -37 | -32 | 12 |
| | | | 2 | posterior | -40.33 | -30.75 | 11.67 |
| **p7** | right | H_1 | 1 | posterior | -45 | -23.36 | -0.09 |
| | | | 2 | posterior | -35.9 | -28.4 | 7 |
| | | NA | 8 | anterior | -39 | -15 | -27 |
| | | OP | 1 | anterior | -30 | -6 | 5 |
| | | | 2 | anterior | -33 | -6 | 6 |
| | | | 3 | anterior | -36 | -5.88 | 7.13 |
| | | | 4 | anterior | -39.18 | -5.45 | 7.91 |
| **p8** | left | OPP_1 | 1 | posterior | 37 | 0.13 | 3.88 |
| | | | 2 | posterior | 41 | 0.67 | 6.89 |
| | | | 3 | posterior | 44.7 | 2 | 7.9 |
| | | OPA | 1 | anterior | 43 | 9 | -15 |
| | | | 2 | anterior | 45 | 9 | -12.86 |
| | | | 3 | anterior | 48 | 9 | -10.67 |
| **p9** | right | OF | 2 | anterior | -37 | 10.14 | -4 |
| | | | 3 | anterior | -40 | 12 | -1 |
| **p10** | left | LOP_1 | 1 | posterior | -31 | -19 | 19 |

| | | | | | | | |
|---|---|---|---|---|---|---|---|
| | | | 2 | posterior | -34 | -19 | 20 |
| | | | 3 | posterior | -37.45 | -19 | 20.09 |
| | | LFT | 1 | anterior | -34 | 29.13 | -3.13 |
| | | | 2 | anterior | -36.27 | 30 | -3.36 |
| | | | 3 | anterior | -38.88 | 30.25 | -4 |
| | | LOF_1 | 1 | anterior | -34 | 25 | 10 |
| | | | 2 | anterior | -36.86 | 26 | 9.43 |
| | | | 3 | anterior | -40.25 | 26.13 | 9 |
| | | LOC | 1 | anterior | -35 | 9 | 4 |
| | | | 2 | anterior | -38.75 | 9 | 4.88 |
| | | | 3 | anterior | -41 | 9 | 5.86 |
| | | | 4 | anterior | -43 | 9 | 7.86 |
| | | LIA_1 | 1 | anterior | -33 | 18 | -7 |
| | | | 2 | anterior | -32 | 18.5 | -4 |
| | | | 3 | anterior | -31 | 19 | -1 |
| | | | 4 | anterior | -29.89 | 19 | 2.44 |
| | | | 5 | anterior | -30 | 17 | 5 |
| | | | 6 | anterior | -29 | 18 | 8 |
| | | | 7 | anterior | -28 | 18 | 12 |
| **p11** | left | INP | 1 | posterior | 42.46 | 0.82 | -16.7 |
| | | | 2 | posterior | 42.49 | -2.29 | -12.89 |
| | | | 3 | posterior | 42.47 | -4 | -9.29 |
| | | | 4 | posterior | 42.52 | -4.88 | -4.32 |
| | | | 5 | posterior | 40.47 | -8.14 | -1.56 |
| | | | 6 | posterior | 40.49 | -10 | 2.4 |
| | | | 7 | posterior | 39.5 | -11.15 | 6.3 |
| | | | 8 | posterior | 39.52 | -12.81 | 8.65 |
| | | | 9 | posterior | 38.46 | -16.09 | 12.68 |
| | | | 10 | posterior | 38.54 | -17.88 | 16.6 |
| | | OF_1 | 1 | anterior | 32.65 | 15.17 | 3.58 |
| | | | 2 | anterior | 38.02 | 17.22 | 3.61 |
| | | | 3 | anterior | 42.48 | 18.04 | 5.57 |
| | | INA_2 | 1 | anterior | 39 | 14.87 | -12.74 |
| | | | 2 | anterior | 39 | 16.15 | -9.35 |
| | | | 3 | anterior | 38 | 18 | -5 |
| | | | 4 | anterior | 37 | 19.88 | -1.65 |
| | | | 5 | anterior | 35.91 | 21.64 | 2.82 |
| | | | 6 | anterior | 33.87 | 23 | 7.5 |
| | | OC | 1 | anterior | 38.1 | 4.9 | 11.38 |
| | | | 2 | anterior | 42.92 | 4.9 | 10.32 |
| | | | 3 | anterior | 48.02 | 5.33 | 11.44 |
| **p12** | left | LIP | 1 | posterior | -35 | 5.03 | -14.78 |
| | | | 2 | posterior | -34.38 | 1.69 | -10.85 |
| | | | 3 | posterior | -34.03 | -1.11 | -7 |
| | | | 4 | posterior | -34 | -4 | -3 |

| | | | | | | |
|---|---|---|---|---|---|---|
| | | 5 | posterior | -33.28 | -6.89 | 2.11 |
| | | 6 | posterior | -32.95 | -9.45 | 6.64 |
| | | 7 | posterior | -33.11 | -12.29 | 10.08 |
| | | 8 | posterior | -32.22 | -16.14 | 13.89 |
| | | 9 | posterior | -32 | -19 | 17.97 |
| | | 10 | posterior | -31.95 | -22 | 22.51 |
| | | 11 | posterior | -31.17 | -24.67 | 27.19 |
| | LIA_2 | 2 | anterior | -32 | 16.88 | -18.29 |
| | | 3 | anterior | -32 | 18.06 | -13.03 |
| | | 4 | anterior | -32.08 | 19.94 | -9.06 |
| | | 5 | anterior | -31 | 22.06 | -5.03 |
| | | 6 | anterior | -31 | 23.7 | -2.11 |
| | | 7 | anterior | -29 | 26.08 | 0.73 |
| | | 8 | anterior | -29 | 28.73 | 4.22 |

**Table S.1.** Description and MNI-coordinates of the electrode contacts used in this investigation, implanted in either the anterior or posterior insula.

| response | modality | condition | location | n | statistic | p.adj |
|---|---|---|---|---|---|---|
| Phase-locked response | thermonociceptive | baseline | anterior | 63 | 1718 | **0.000** |
| | thermonociceptive | distraction | anterior | 63 | 1247 | 0.051 |
| | vibrotactile | baseline | anterior | 48 | 999 | **0.000** |
| | vibrotactile | distraction | anterior | 48 | 896 | **0.002** |
| | thermonociceptive | baseline | posterior | 37 | 701 | **0.000** |
| | thermonociceptive | distraction | posterior | 37 | 701 | **0.000** |
| | vibrotactile | baseline | posterior | 24 | 256 | **0.002** |
| | vibrotactile | distraction | posterior | 24 | 285 | **0.000** |
| Theta frequency band | thermonociceptive | baseline | anterior | 62 | 963 | 0.539 |
| | thermonociceptive | distraction | anterior | 63 | 1255 | 0.091 |
| | vibrotactile | baseline | anterior | 48 | 910 | **0.001** |
| | vibrotactile | distraction | anterior | 48 | 1015 | **0.000** |
| | thermonociceptive | baseline | posterior | 37 | 683 | **0.000** |
| | thermonociceptive | distraction | posterior | 37 | 655 | **0.000** |
| | vibrotactile | baseline | posterior | 24 | 252 | **0.004** |
| | vibrotactile | distraction | posterior | 24 | 268 | **0.001** |
| Alpha frequency band | thermonociceptive | baseline | anterior | 63 | 1153 | 0.161 |
| | thermonociceptive | distraction | anterior | 63 | 1269 | 0.075 |
| | vibrotactile | baseline | anterior | 48 | 798 | **0.046** |
| | vibrotactile | distraction | anterior | 48 | 1021 | **0.000** |
| | thermonociceptive | baseline | posterior | 36 | 634 | **0.000** |
| | thermonociceptive | distraction | posterior | 37 | 639 | **0.000** |
| | vibrotactile | baseline | posterior | 24 | 269 | **0.001** |
| | vibrotactile | distraction | posterior | 23 | 247 | **0.001** |
| Beta frequency band | thermonociceptive | baseline | anterior | 63 | 1152 | 0.326 |
| | thermonociceptive | distraction | anterior | 63 | 1258 | 0.131 |
| | vibrotactile | baseline | anterior | 48 | 570 | 0.575 |
| | vibrotactile | distraction | anterior | 48 | 871 | **0.008** |
| | thermonociceptive | baseline | posterior | 37 | 569 | **0.002** |
| | thermonociceptive | distraction | posterior | 37 | 617 | **0.000** |
| | vibrotactile | baseline | posterior | 24 | 218 | 0.105 |
| | vibrotactile | distraction | posterior | 24 | 262 | **0.002** |
| Gamma frequency band | thermonociceptive | baseline | anterior | 63 | 1358 | **0.033** |
| | thermonociceptive | distraction | anterior | 63 | 1126 | 0.346 |
| | vibrotactile | baseline | anterior | 48 | 681 | 0.346 |
| | vibrotactile | distraction | anterior | 48 | 905 | **0.003** |
| | thermonociceptive | baseline | posterior | 37 | 583 | **0.001** |
| | thermonociceptive | distraction | posterior | 37 | 554 | **0.005** |
| | vibrotactile | baseline | posterior | 24 | 243 | **0.016** |
| | vibrotactile | distraction | posterior | 24 | 221 | 0.064 |

**Table S.2.** Results of the right-tailed Wilcoxon signed-rank test against 0 to assess whether the amplitude at the FOI was significantly different from zero for each modality, condition and electrode contact location. Holm-correction was applied to correct for multiple comparisons.

| | contrast | location | estimate | SE | df | lower.CL | upper.CL | t | p |
|---|---|---|---|---|---|---|---|---|---|
| Phase-locked | thermonociceptive - vibrotactile | anterior | -0.00 | 0.22 | 269.45 | -0.43 | 0.43 | -0.01 | 1.00 |
| | thermonociceptive - vibrotactile | posterior | 1.89 | 0.31 | 284.51 | 1.29 | 2.49 | 6.18 | **0.00** |
| Theta | thermonociceptive - vibrotactile | anterior | -0.26 | 0.05 | 265.30 | -0.36 | -0.16 | -4.99 | **0.00** |
| | thermonociceptive - vibrotactile | posterior | 0.16 | 0.07 | 277.93 | 0.02 | 0.30 | 2.28 | **0.02** |
| Alpha | thermonociceptive - vibrotactile | anterior | -0.09 | 0.02 | 256.90 | -0.13 | -0.04 | -3.56 | **0.00** |
| | thermonociceptive - vibrotactile | posterior | 0.10 | 0.03 | 268.68 | 0.03 | 0.17 | 2.91 | **0.00** |
| Gamma | thermonociceptive - vibrotactile | anterior | -0.01 | 0.01 | 267.69 | -0.02 | 0.01 | -0.90 | 0.37 |
| | thermonociceptive - vibrotactile | posterior | 0.03 | 0.01 | 283.58 | 0.00 | 0.05 | 2.37 | **0.02** |

**Table S.3.** Post-hoc pairwise comparisons of model estimated marginal means based on significant interactions *modality* x *location* in the LMM of each frequency band. Significant comparisons ($p<0.05$) are indicated in bold font. These results are illustrated in Figure 3.

| | modality | condition | location | n | statistic | p.adj | d |
|---|---|---|---|---|---|---|---|
| Phase-locked response | **thermonociceptive** | **baseline** | **anterior** | **63** | **4.808** | **0.000** | **0.606** |
| | thermonociceptive | baseline | posterior | 37 | -0.521 | 1.000 | -0.086 |
| | vibrotactile | baseline | anterior | 48 | -0.496 | 1.000 | -0.072 |
| | vibrotactile | baseline | posterior | 24 | 0.124 | 1.000 | 0.025 |
| Theta frequency band | thermonociceptive | baseline | anterior | 62 | -1.266 | 0.528 | -0.161 |
| | thermonociceptive | baseline | posterior | 37 | 0.106 | 0.917 | 0.017 |
| | **vibrotactile** | **baseline** | **anterior** | **48** | **-3.316** | **0.007** | **-0.479** |
| | vibrotactile | baseline | posterior | 24 | -1.397 | 0.528 | -0.285 |
| Alpha frequency band | thermonociceptive | baseline | anterior | 63 | -0.257 | 0.968 | -0.032 |
| | thermonociceptive | baseline | posterior | 36 | 1.035 | 0.924 | 0.172 |
| | **vibrotactile** | **baseline** | **anterior** | **48** | **-4.077** | **0.001** | **-0.589** |
| | vibrotactile | baseline | posterior | 23 | -0.712 | 0.968 | -0.148 |
| Beta frequency band | thermonociceptive | baseline | anterior | 63 | -0.267 | 1.000 | -0.034 |
| | thermonociceptive | baseline | posterior | 37 | 0.426 | 1.000 | 0.070 |
| | vibrotactile | baseline | anterior | 48 | -2.494 | 0.065 | -0.360 |
| | vibrotactile | baseline | posterior | 24 | -2.166 | 0.123 | -0.442 |
| Gamma frequency band | thermonociceptive | baseline | anterior | 63 | 1.785 | 0.238 | 0.225 |
| | thermonociceptive | baseline | posterior | 37 | -1.064 | 0.588 | -0.175 |
| | **vibrotactile** | **baseline** | **anterior** | **48** | **-2.680** | **0.040** | **-0.387** |
| | vibrotactile | baseline | posterior | 22 | -0.053 | 0.958 | -0.011 |

**Table S.4.** A one-sample t-test with a Holm-correction assessing whether the relative amplitude ($\Delta$ amplitude = baseline - distraction) is significantly different from 0, thereby indicating a difference between the conditions irrespective of the magnitude of the amplitude at baseline. Significant comparisons ($p<0.05$) are indicated in bold font. These results are also illustrated in Figure 5.
